# Overconfidence undermines global wildlife abundance trends

**DOI:** 10.1101/2022.11.02.514877

**Authors:** Thomas Frederick Johnson, Andrew P Beckerman, Dylan Z Childs, Christopher A Griffiths, Pol Capdevila, Christopher F Clements, Marc Besson, Richard D. Gregory, Eva Delmas, Gavin Thomas, Karl Evans, Tom Webb, Rob Freckleton

**Affiliations:** School of Biosciences, University of Sheffield, Western Bank, S10 2TN, Sheffield, UK; Swedish University of Agricultural Sciences, Department of Aquatic Resources, Institute of Marine Research, Turistgatan 5, SE-453 30, Lysekil, Sweden; School of Biological Sciences, Biosciences, University of Bristol, 24 Tyndall Ave, BS8 1TQ, BristolSheffield, Western Bank, S10 2TN, Sheffield, UK; Departament de Biologia Evolutiva, Ecologia i Ciències Ambientals, Universitat de Barcelona, Avda. Diagonal 643, 08028 Barcelona; School of Biological Sciences, University of Bristol, Life Sciences Building, 24 Tyndall Avenue, Bristol, BS8 1TQ, UK; School of Biological Sciences, University of Bristol, 24 Tyndall Ave, BS8 1TQ, Bristol, UK; Sorbonne Université CNRS UMR Biologie des organismes marins, BIOM, F-66650 Banyuls-sur-Mer, France; RSPB Centre for Conservation Science, The Lodge, Sandy, Bedfordshire SG19 2DL, UK; Centre for Biodiversity & Environment Research, Department of Genetics, Evolution and Environment, University College London, Darwin Building, Gower Street, London, WC1E 6BT, UK; Habitat, 5818 Boulevard Saint-Laurent, Montréal (QC), H2T 1T3, Canada; Institut des Sciences de la Forêt Tempérée, Université du Québec en Outaouais, 58 rue Principale, Ripon (QC) J0V 1V0, Canada

## Abstract

In the face of rapid global change and an uncertain fate for biodiversity, it is vital to quantify trends in wild populations. These trends are typically estimated from abundance time series for suites of species across large geographic and temporal scales. Such data implicitly contain phylogenetic, spatial, and temporal structure which, if not properly accounted for, may obscure the true magnitude and direction of biodiversity change. Here, using a novel statistical framework to simultaneously account for all three of these structures, we show that the majority of current abundance trends estimates among 10 high-profile datasets, representing millions of abundance observations, are likely unreliable or incorrect. Our new approach suggests that previous models are too simplistic, incorrectly estimating global abundance trends and often dramatically underestimating uncertainty, an aspect that is critical when translating global assessments into policy outcomes. Further, our approach also results in substantial improvements in abundance forecasting accuracy. Whilst our results do not improve the outlook for biodiversity, our framework does allow us to make more robust estimates of global wildlife abundance trends, which is critical for developing policy to protect our biosphere.

## Main

Accelerating rates of species extinction are driving global changes in biodiversity, threatening ecosystems and the services they provide (*1*). In an attempt to reverse biodiversity declines, world leaders, policymakers, and academics have called for action (*2*). Evidence-based actions require long-term datasets and rigorous modelling to produce metrics that can reliably describe biodiversity change through time (*3*). Currently, some of the most influential metrics are calculated using datasets like BioTIME (*4*), the Living Planet (*5*), or the North American Breeding Bird Survey (*6*), which shape our understanding of global wildlife abundance trends. The data that underpin these metrics – and thus policy decisions – are time series of wildlife population abundances derived from long-term monitoring schemes conducted across multiple years, species, and locations. Such data currently represent a key pillar of global biodiversity monitoring but are potentially prone to a series of data biases which complicate the statistical interpretation of the current and historic wildlife abundance trends (*7*–*9*).

A formidable challenge in reliably estimating abundance trends is non-independence in the data (*10*). Non-independence may present in a variety of ways, which we split into two core types: *Hierarchical* - where observations are pseudoreplicated e.g. multiple trends for a given species; and *Correlative* - where observations become increasingly correlated (sometimes termed autocorrelation) when close in space (*11*), time (*12*), or phylogeny (*13*), such that we may expect more closely related species to have more similar trends - Figure S1. Whilst studies commonly account for hierarchical non-independence using features like random effects in mixed models (Table S2), no method has yet been formalised to account for all three sources of correlative non-independence simultaneously.

Here we introduce a comprehensive statistical framework, the *correlated effect* model, that controls for both hierarchical non-independence and all three sources of correlative non-independence, and apply it to 10 high-profile, multi-species datasets (Table S1) which have been used to infer abundance trends in global biodiversity (*4*, *6*, *14*–*21*). Combined, these datasets describe the abundance patterns of more than 27,000 populations, representing ~1,700 species and ~2,800 unique locations, and are considered some of the best biodiversity monitoring datasets available. We compare our correlated effect model with two mixed-effect modelling frameworks (random intercept and random slope) that only account for hierarchical non-independence and are commonly used in the literature (Figure 1). All three models have the same core structure and purpose - defining abundance as a function of time for each population, clustered into species and location hierarchies, which are then averaged within the model to derive a ‘global abundance trend’. We show that the commonly-used approaches that ignore correlative non-independence often misrepresent trends, by misestimating trend direction (e.g. negative trends become positive) and underestimating uncertainty (e.g. trend variance inflates substantially).

**Figure 1.**
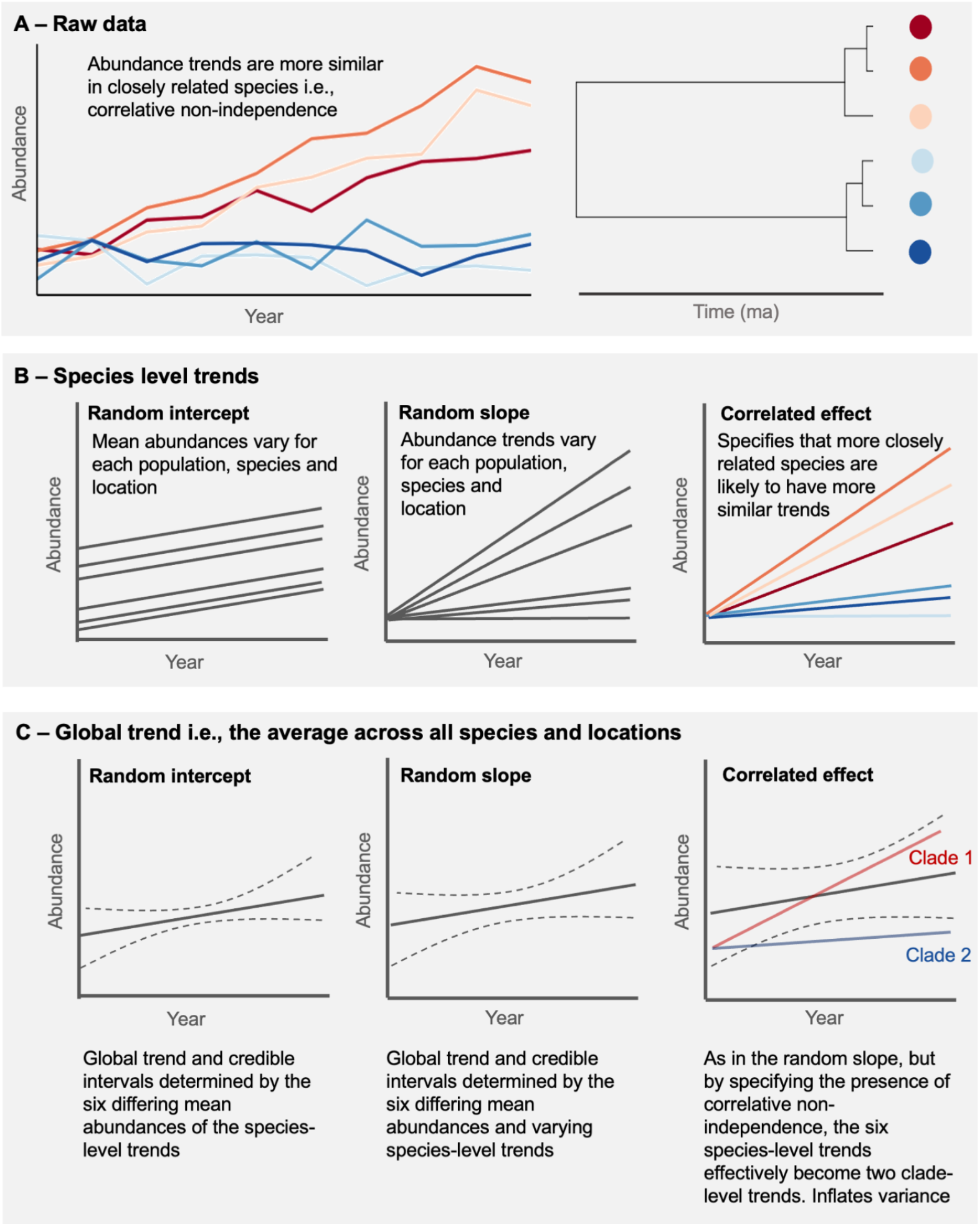
Proposed impact of correlative non-independence on global abundance trends. A) Abundance time-series for six species in a given location, simulated to exhibit strong phylogenetic signal in their abundance-year slope coefficient i.e. abundance changes are more similar in more closely related species. Time-series colour corresponds with the adjacent phylogeny where species with more similar colours are more closely related. B) Approach of our three models for estimating these species level abundance trends, notably, the correlated effect model differentiates from the random slope model as it acknowledges the potential presence of correlative non-independence i.e. phylogenetic signal in species trends. C) Estimates of global abundance trends and credible intervals (i.e. the overall fixed effect abundance-time coefficient and credible intervals across all species and locations) for each of the three models - depicting the proposed impact of correlative non-independence (phylogenetic signal in this example) on global trend estimates. This figure purely represents phylogenetic correlative non-independence, but the correlated effect model also captures spatial (i.e. trends are likely more similar when close in space) and temporal (abundance observations are likely more similar when close in time) correlative non-independence.

## Accounting for correlative non-independence changes global abundance trends

Across the 10 datasets and relative to the variance captured by all residual, fixed and random effects, the temporal (mean = 0.29, sd = 0.19), spatial (mean = 0.13, sd = 0.09), and phylogenetic (mean = 0.12, sd = 0.08) components accounted for a substantial proportion of variance in the correlated effect model. The correlated terms accounted for nearly half of this variance with a mean *h -* variance captured by the correlated terms divided by the combined variance of the correlated and hierarchical terms *-* of 0.45 (sd = 0.15) for the spatial terms and 0.49 (sd = 0.23) for the phylogenetic terms. Covariance between neighbouring abundance observations through time was very high with a *rho -* the correlation between neighbouring abundance observations *-* of 0.77 (sd = 0.30). These results provide evidence that abundance-trends exhibit temporal, spatial, and phylogenetic patterns.

Global abundance trends (i.e. the average abundance trend across all species and locations) change substantially when correlative non-independence is accounted for (Fig 2). We find that the correlated effect model reverses trends (e.g., goes from -ve to +ve) in 5 of the 10 datasets, relative to the random intercept model, and 3 out of 10 datasets relative to the random slope model.

**Figure 2.**
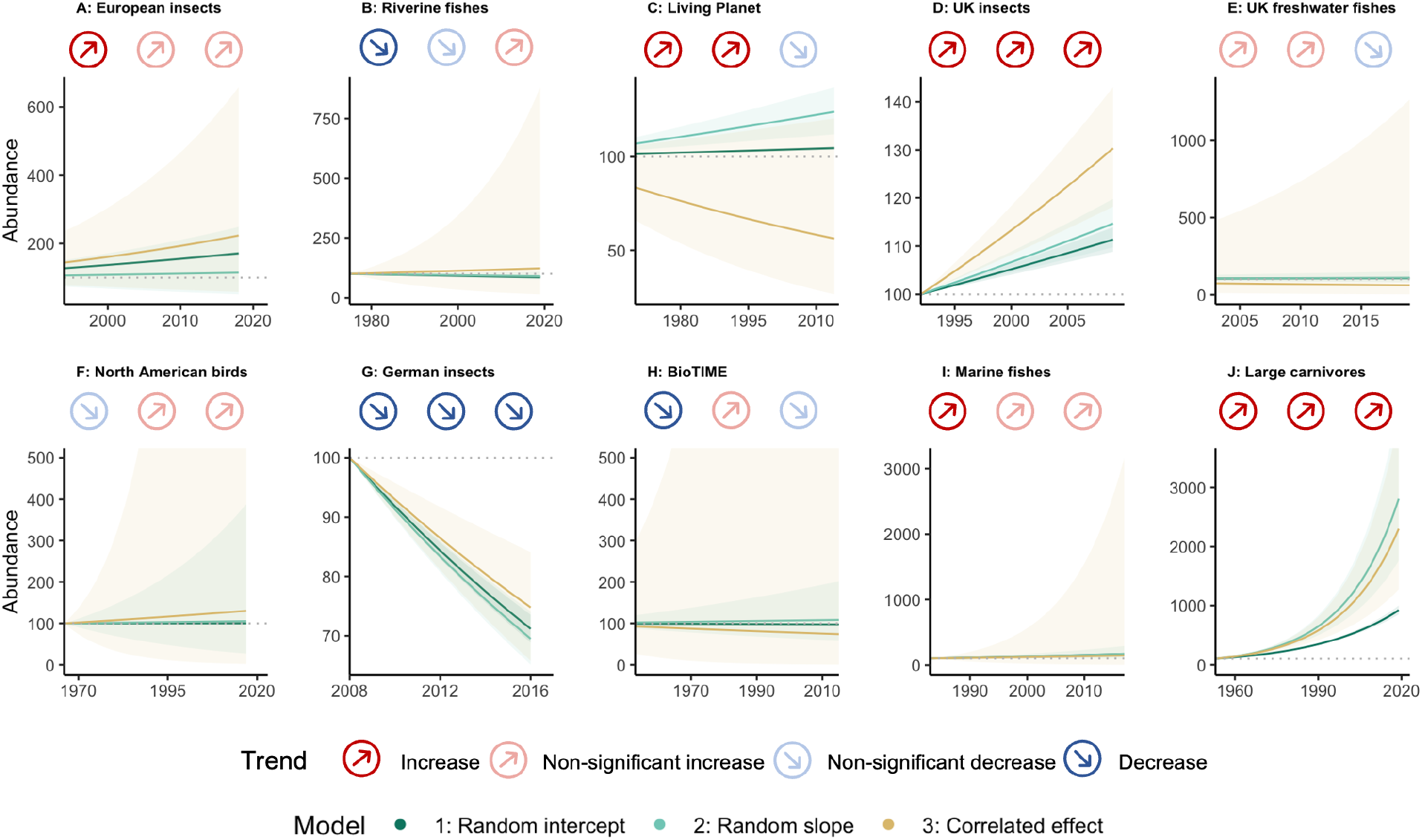
Widely-used statistical models misrepresent biodiversity abundance trends. Abundance trends and projections across 10 high-profile datasets under three different models. Trends (circled arrows), describe the global change in abundance (averaged across all species and locations) for each dataset in our three models (from left to right): random intercept, random slope, and the correlated effect model which simultaneously accounts for three different sources of correlative non-independence in time, space and phylogeny. We specify four categories of trend: Increase – median abundance-time coefficient is positive and significant at the 50% credible interval; Non-significant increase – median abundance-time coefficient is positive, but not significant at the 50% credible interval; Non-significant decrease – median abundance-time coefficient is negative, but not significant at the 50% credible interval; Decrease – median abundance-time coefficient is negative and significant at the 50% credible interval. We use the global trend coefficient (average across all species and locations) and 50% credible intervals (represented by shading) to produce abundance projections for each model in each dataset from an arbitrary baseline abundance of 100. Projections cover the time-span of the observed data.

Global abundance trend uncertainty (i.e. the variance around the global abundance-time coefficient) was under-estimated in all 10 datasets in both the random intercept and random slope models. These under-estimates are dramatic, with uncertainty in the correlated effect model being 70 times greater [95% CI: 29 - 172] than the random intercept model and 4 times greater [95% CI: 2 – 10] than the random slope model (Figure S2). The degree of change between the 3 models is determined by the presence and amount of correlative non-independence, with high signal (h) positively associated with inflation in uncertainty (Figure S3).

The implications of our findings for global estimates of biodiversity are substantial. Firstly, the global trends estimated from these policy-shaping datasets often change after accounting for correlative non-independence (Figure 2). Across the three models, there is only agreement (trends have the same direction and significance status) in 2 of the datasets. For instance, when using the Living Planet data, which has been used to determine the conservation status of vertebrate populations worldwide (*5*), we estimate a reversal from a significant increase (at 50% credible intervals) in the random intercept and random slope models, to a non-significant decline in the correlated effect model. Notably, our Living Planet result differs from the Living Planet Index, which uses an entirely different modelling method which we do not test (see Table S2), and a larger sample of data (we use the longest 50% of time series in each dataset).

After accounting for correlative non-independence, global abundance trends are unclear, with error bars overlapping zero at the 50% credible interval (indicative of no clear trend) in eight of the 10 datasets. In contrast, only two and six datasets overlap zero in the random intercept and random slope models, respectively. This discrepancy is driven by an under-estimation of variance by the simpler models such that estimated declines and recoveries in the random intercept and slope models disappear when correlative non-independence is appropriately accounted for. Further, using the more commonly-applied 95% credible intervals, the correlated effect model overlaps zero in all but one dataset, suggesting that abundance trends and their associated conclusions on biodiversity change are uncertain across the majority of our high-profile datasets.

## Accounting for non-independence improves prediction

Despite the large uncertainty among the datasets, accounting for correlative non-independence improves our capacity to make predictions ‘out of sample’. Global-scale abundance trends are vital to policy and have fed into the IPBES assessment (*1*) and Aichi targets (*22*); part of the value of these abundance trends is predicated on being able to estimate which species and locations are likely to be declining or recovering, and when. One of the benefits of accounting for correlative non-independence is that it has the potential to not only improve our estimation of global trends, but also improves our ability to predict outside the scope of the data. Using the largest abundance dataset – BioTIME (Table S1) - as a case study, we tested each model’s ability to forecast new abundance observations and estimate population trends. This involved removing the final abundance observation in 50% (n = 3520) of the population abundance time series and then evaluating each of the three models’ ability to predict this value. Next, we removed all abundance observations for 10% (n = 704) of populations and assessed each of the three models’ ability to use the species and location information to predict the removed population trends i.e. can we estimate trends in populations without data? We report predictive accuracy using the median absolute error (*MAE*) in the predicted value minus the observed value, where a larger error indicates worse predictive performance.

The correlated effect model was able to estimate the final abundance observation with only a 5% error (MAE = 0.05), three times more accurately than the random intercept model (MAE = 0.15) and 2.4 times more accurate than the random slope models (MAE = 0.12; Fig S4). The correlated effect also performed best when estimating missing population trends, with an error of only 1.34%, marginally better than the 1.39% of the random slope model (Fig S5) – we did not compare the random intercept model which would specify that all populations have the same trend.

The improved prediction of abundances by the correlated effect model is a consequence of handling the temporal non-independence in BioTIME (*rho* = 0.56). This correlation between neighbouring abundance values introduces nonlinearity (residual variability in linear trends) in temporal trends, and more closely represents realistic population dynamics (Fig 3: *Population-level*). At coarser ecological scales, handling sources of correlative non-independence could also influence species-level and site-level trends. For example, handling correlative non-independence increases the variance in the site-level trend (i.e. the trend for a given location) relative to the random slope model (Fig 3: *Site-level*). At the global-level, the presence of temporal, spatial (*h* = 0.59), and phylogenetic (*h* = 0.72) signals inflates the uncertainty around the overall trend and leads to shifts in the mean trend (Fig S3). However, the presence of this signal also has the added benefit of allowing us to make predictions beyond the species and location data (Fig 4), offering important insight into species and locations most likely to exhibit declines and recoveries.

**Figure 3.**
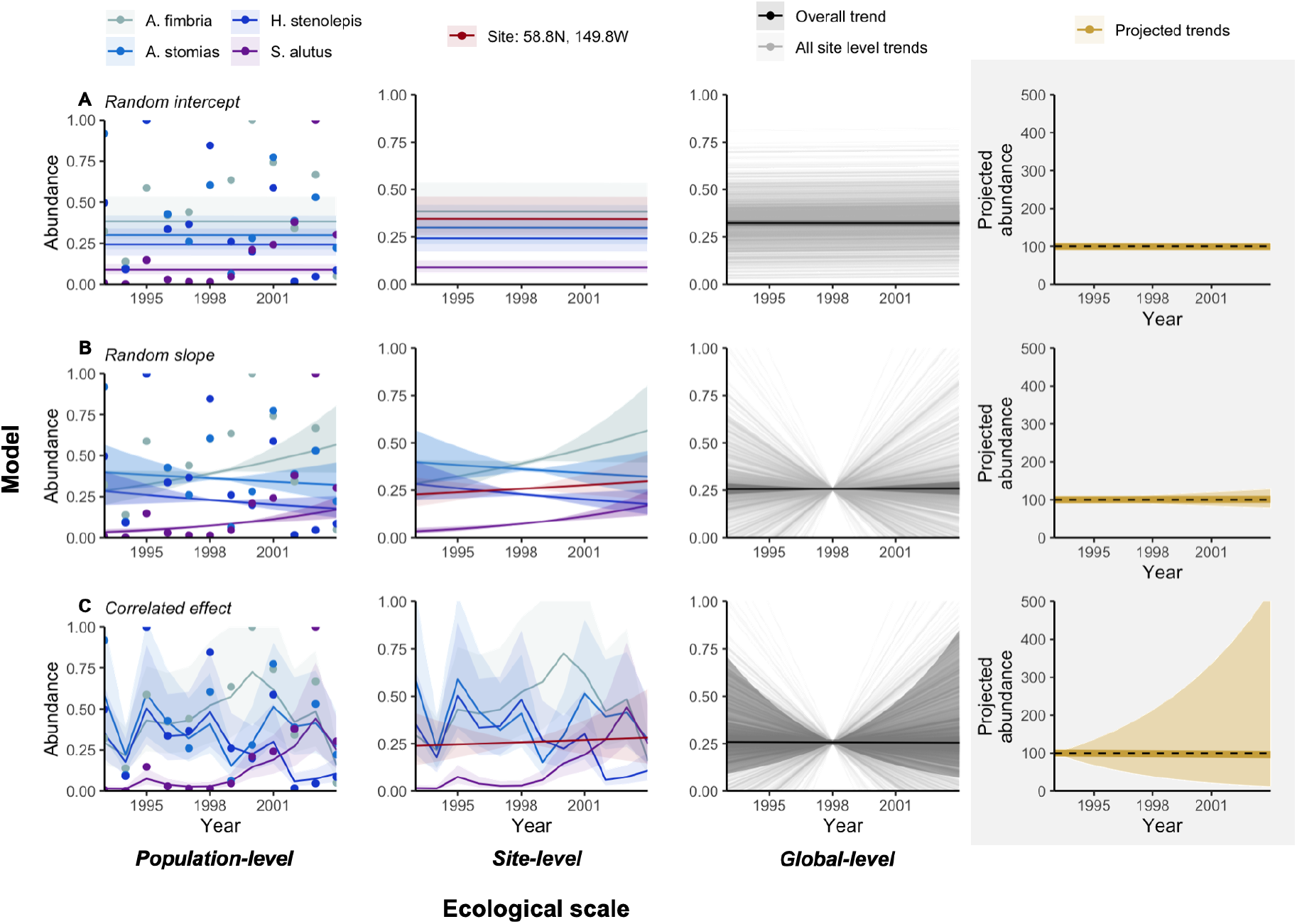
More complex models better represent population dynamics and improve the validity of conclusions across ecological scales. Example of how the three models (each on a row; A: Random intercept, B: Random slope, and C: Correlated effect) describe abundance patterns at different ecological scales (finer ecological scales on the left). The population-level column showcases how each of the three models produce different estimates of abundance trends (lines and 95% credible interval shading) for all four species with data in this given location, with data points representing the observed abundance values. The location-level column depicts how the species’ trends, under different models, influence the site-level trend (i.e. a trend for a given location; red), where the line and 95% credible intervals describe the average trend and variability in trend (respectively) across all species in the given location. At the global-level, the trend for each unique site is represented by a faded-grey line, and the global trend coefficient and 95% credible intervals are depicted by the black line and shading. The grey-box in the final column describes how a hypothetical population would change under the BioTIME global trend coefficient and 95% credible intervals projected from a relative baseline abundance of 100. This example is based on data within BioTIME. In each plot, we restrict the time-frame to the temporal extent of the population-level trends (1993 – 2004), instead of the total temporal extent of our BioTIME sample.

**Figure 4.**
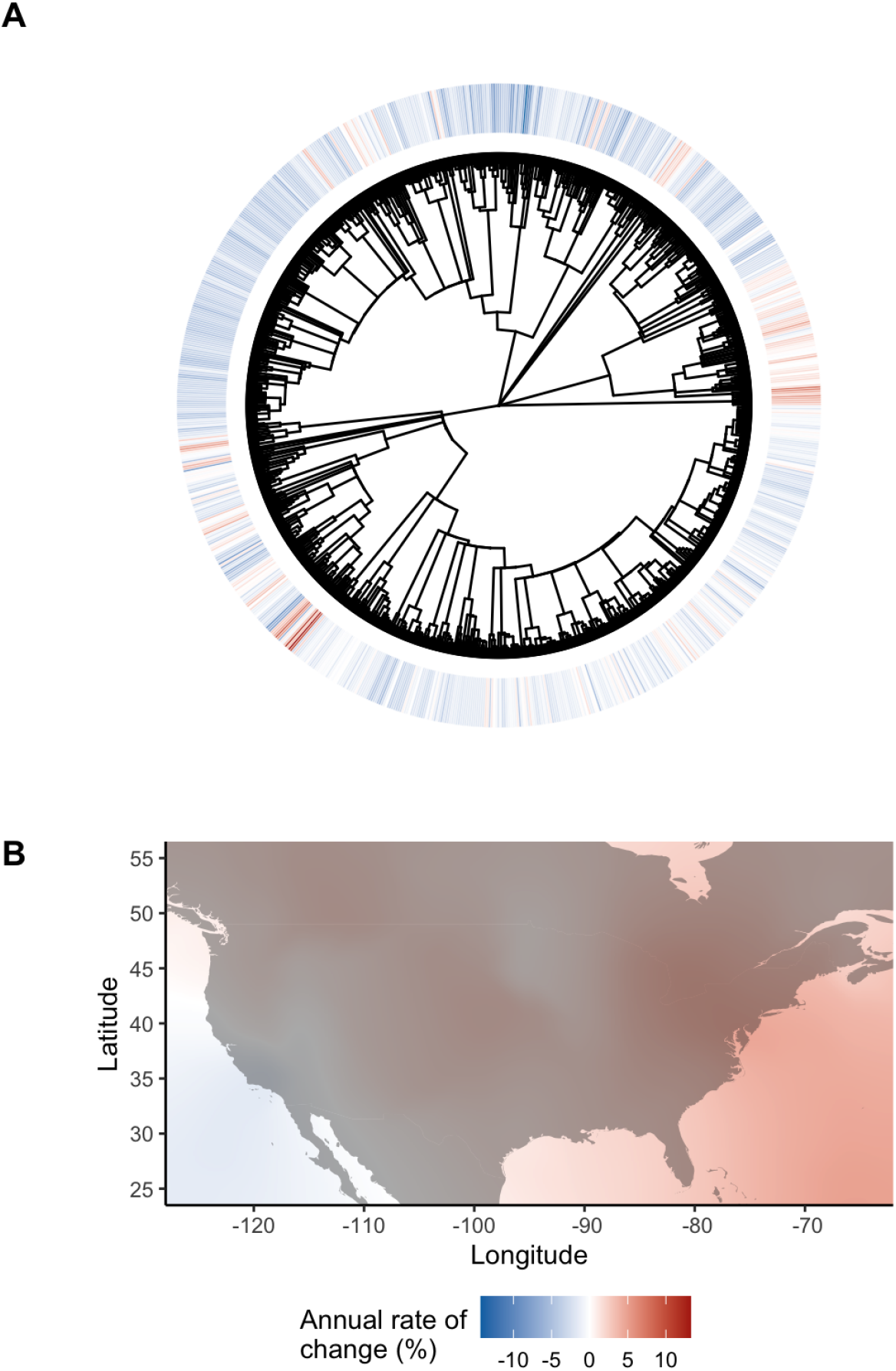
Phylogenetic and spatial distribution of abundance change. Median predicted abundance change across a phylogeny and space. In the phylogeny (A), the median change describes the estimated annual rate of change (%) in abundance for each species averaged across all sites (locations) and populations, whilst over space (B), the median change describes estimates for each location averaged over all species and populations. In both panels, the median change is drawn solely from the phylogenetic and spatial correlation terms in the correlated effect model e.g., for the phylogeny (A), change describes solely the phylogenetic signal. For the phylogeny, we have log-scaled the branch lengths. For space (B), abundance change has been down-scaled from a 5° to 0.1° resolution using interpolation. These data are drawn from BioTIME (Table S1).

## Implications for biodiversity change

The abundance datasets we analyse are highly influential within policy, and so it is vital that any inference gained is both valid and reliable. Our work shows that when biodiversity change datasets are analysed without accounting for correlative non-independence, and only with hierarchical structure, there is a substantial risk that global average population trends could be mis-estimated and that too much confidence is placed on these estimates. Use of the existing methods therefore creates a perfect-storm where increases may be reported as declines, and vice versa, and these conclusions are made with unjustified confidence. This raises serious questions about the validity of the existing methods for describing global or regional abundance change where datasets are spatially, temporally, or phylogenetically non-independent (Table S2). Including correlated effects to capture well known issues of space, time and phylogeny addresses this issue in two critical ways: 1) improving our global trend estimate, and 2) highlights the enormous uncertainty associated with this estimate - answering calls for more fair representation of uncertainty in ecological studies (*23*). In fact, nine out of 10 of our global-trends are not significant at conventional 95% credible interval thresholds, so whilst populations may well be declining in specific species or locations, we find little evidence of systematic abundance declines (*7*, *24*). These results do not necessarily mean that wildlife abundances have not declined, or that the current estimates of trends are incorrect. It is possible that abundances may have increased on average, or perhaps declines have been far more extreme than we have previously imagined; simply, the uncertainty is too high to know.

Whilst the high uncertainty around trends limits our understanding of global-scale abundance changes, the improved prediction from the correlated effect model offers a pathway towards estimating species and locations where populations are likely to be declining (Figure 4). This is a key advancement and may help inform species extinction risk. However, given the severity of the potential implications of abundance declines (*25*), it is vital that we urgently expand our methods, utilising modern modelling approaches (*26*) and data collection philosophies (*27*) to help answer the ultimate question for biodiversity change researchers with some precision - how is biodiversity changing?

## Acknowledgements

Thanks to all who have provided open-data or enabled access to data: Verstraeten A., Neirynck J., Van Ryckegem G., Van Hoey G., Nikolov B., Evtimova V., Stanchev R., Haase P., Kronck I., Meyer J., Creveceour L., Janssen M., Thimonier A., Pilotto F., Kappel Schmidt I., Oro D., Poyry J., Huttunen K., Muotka T., Paavola R., Barboro L., Camatti E., Pansera M., Alber R., Vorhauser S., Skula A., Springe G., Ens B.J., Visser M. E., Bongard T., Jensen T., Baekkelie K. A. E., Jensen T. C., Pardal M., Martinho F., Pipan T., Aljancic M., Gabrovsek F., Kalivoda H., Halada L., David S., Grandin U., Monteith D., Rennie S., Adamson J., Anderson R., Andrews C., Bater J., Bayfield N., Beaton K., Beaumont D., Benham S., Bowmaker V., Britt C., Brooker R., Brooks D., Brunt J., Common G., Cooper R., Corbett S., Critchley N., Dennis P., Dick J., Dodd B., Dodd N., Donovan N., Easter J., Flexen M., Gardiner A., Hamilton D., Hargreaves P., Hatton-Ellis M., Howe M., Kahl J., Lane M., Langan S., Lloyd D., McCarney B., McElarney Y., McKenna C., McMillan S., Milne F., Milne L., Morecroft M., Murphy M., Nelson A., Nicholson H., Pallett D., Parry D., Pearce I., Pozsgai G., Rose R., Schafer S., Scott T., Sherrin L., Shortall C., Smith R., Smith, P., Tait R., Taylor C., Taylor M., Thurlow M., Tilbury C., Turner A., Tyson K., Watson H., Whittaker M., Wood C., Winfield I., Di Fonzo M.I., Collen, B., Chauvenet A.L.M., Mace G.M., Comte L., Carvajal Q.J., Tedesco P.A., Giam X., Brose U., Erős T., Filipe A.F., Fortin M.J., Irving K., Jacquet C. and Larsen S., Sauer J. R., Niven D. K., Hines J. E., Ziolkowski D. J., Pardieck Jr. K. L., Fallon J. E., Link W. A., Thackeray, S.J., Dornelas M., Antão L.H., Moyes F., Bates A.E., Magurran A.E., Greenstreet S.P.R, Moriarty M., Batten S.D., Seibold S., Weisser W., Gossner M., Pasalic E., Lange M., Turke M., Gallenberger S.N., and Staab M. Also, thanks to everyone contributing data to open databases, including: BioTIME, the Living Planet Index, UK Environment Agency.

