## Supplementary material for "Overconfidence undermines global wildlife abundance trends"

#### Materials and methods

Data

To test how accounting for the presence of both hierarchical and correlative non-independence (Figure S1) impacts inference, we compiled 10 datasets that describe population abundances through time. These datasets represent some of the most influential within ecology and conservation biology, forming a basis for policy-shaping reports like the Living Planet (*1*) and IPBES (*2*), as well as a series of high-profile and highly-cited publications - see Table S1 for a full list. For each dataset, we extracted the population abundance estimates, the accompanying time-stamps, the species scientific names, the name of the site (location) where the population was sampled, and any site coordinates (where possible). Whilst these datasets are vital within biodiversity science, many of the dataset are prone to biases e.g. lacking tropical representation, and contain few plant and invertebrate species.

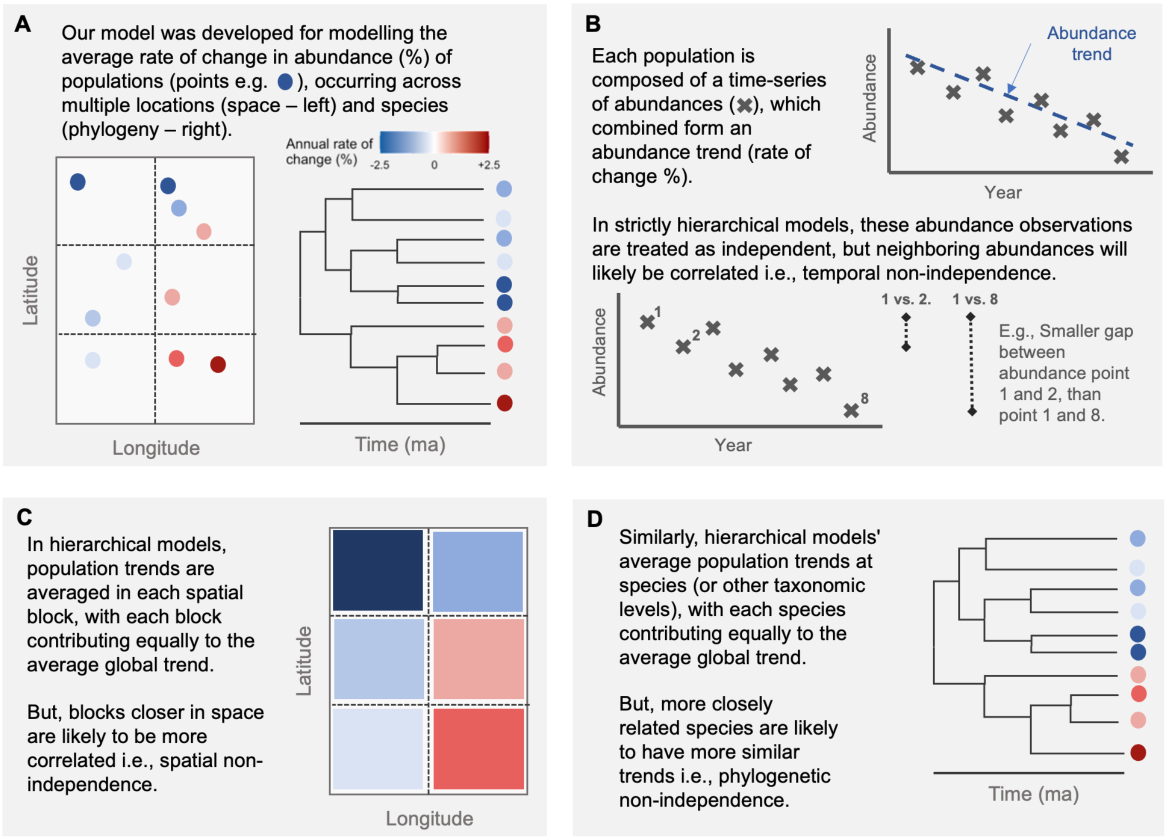

**Figure S1.** Conceptual infographic depicting how both hierarchical and correlative non-independence could occur and be accounted for in large abundance datasets.

**Table S1.** Dataset name and description for each of our 10 datasets. Descriptions include temporal, taxonomic and spatial summaries.

| Dataset | Description |
|---|---|
| A: European insects (*3*) | Population abundance time series for European insects. Covering 79 abundance time series, derived from 2,819 abundance observations. These time series represent 7 unique sites and 61 species.<br><br>Temporal extent: 1974 - 2018<br>Latitude extent: 48.7 – 65.0<br>Longitude extent: -0.9 - 24.6<br><br>Notes: We extracted relative abundance/density estimates from across 51 datasets used by Pilotto et al (2020). We excluded a further 31 datasets utilised by Pilotto et al (2020) as the data lacked clear metadata, or data were not resolved to the species level, or data represented species presence/absences instead of abundances. |
| B: Riverine fishes(*4*) | Population abundance time series for riverine fishes. Covering 679 abundance time series, derived from 17,435 abundance observations. These time series represent 120 unique sites and 83 species.<br><br>Temporal extent: 1975 - 2018<br>Latitude extent: 48.7 – 65.0<br>Longitude extent: -0.9 - 24.6 |
| C: Living Planet (*5*) | Global population abundance time series for vertebrates. Covering 709 abundance time series, derived from 27,689 abundance observations. These time series represent 348 unique sites and 487 species.<br><br>Temporal extent: 1950 - 2014<br>Latitude extent: -77.8 - 74.8<br>Longitude extent: -176.0 - 173.5<br><br>Notes: Used a 2016 batch release so abundance data from recent years are excluded |
| D: UK insects (*6*) | Population abundance time series for UK insects. Covering 146 abundance time series, derived from 2,710 abundance observations. These time series represent 1 unique site and 146 species.<br><br>Temporal extent: 1992-2009<br><br>Notes: Available data lack spatial information |

| | |
|---|---|
| E: UK freshwater fishes (*7*) | Population abundance time series for UK freshwater fishes. Covering 491 abundance time series, derived from 4,036 abundance observations. These time series represent 208 unique sites and 21 species.<br><br>Temporal extent: 1981-2020<br>Latitude extent: 50.9 - 55.1<br>Longitude extent: -3.6 - 1.6 |
| F: North American Breeding Birds (*8*) | Population abundance time series from the North American breeding bird survey. Covering 4,059 abundance time series, derived from 211,116 abundance observations. These time series represent 29 unique sites and 260 species.<br><br>Temporal extent: 1966-2017<br>Latitude extent: 28.6 - 46.8<br>Longitude extent: -90.0 - -63.4<br><br>Notes: Available data are summarised at the state/province level for the US and Canada |
| G: German insects (*9*) | Population abundance time series for German insects. Covering 160 abundance time series, derived from 1,531 abundance observations. These time series represent 1 unique site and 43 species.<br><br>Temporal extent: 2008-2016<br><br>Notes: Available data lack spatial information. Data that were not resolved to the species level were excluded. |
| H: BioTIME (*10*) | Population abundance time series from the BioTIME dataset - representing all core taxa and realms. Covering 6,511 abundance time series, derived from 194,451 abundance observations. These time series represent 291 unique sites and 316 species.<br><br>Temporal extent: 1953-2015<br>Latitude extent: 26.3 - 66.0<br>Longitude extent: -123.6 - 16.3 |
| I: Marine fishes (*11*) | Population abundance time series for marine fishes in the North Sea. Covering 620 abundance time series, derived from 21,765 abundance observations. These time series represent 123 unique sites and 25 species.<br><br>Temporal extent: 1983-2017<br>Latitude extent: 51.1 - 61.2<br>Longitude extent: 0.4 - 12.3 |
| J: Large carnivores (*12*) | Population abundance time series for large carnivores. Covering 76 abundance time series, derived from 1,427 abundance observations. These time series represent 76 unique sites and |

| | |
|---|---|
| | 17 species.<br><br>Temporal extent: 1953 - 2019<br>Latitude extent: -28 - 67<br>Longitude extent: -143 - 39 |

Using the species scientific names, we searched and extracted a synthetic tree from the open tree of life (*13*, *14*). The open tree of life is a database resulting from the collective effort of pooling existing phylogenetic trees and taxonomies into a shared topology. In recent years, it has been used to explore host-symbiont interactions (*15*), drivers behind fish growth (*16*), and the impacts of climate change on plants and herbivores (*17*). This topology lacks branch lengths (the evolutionary distance between nodes), which we estimated using Grafen's approach (*18*) from the compute.brlen function in the R package ape (*19*). The lack of branch lengths and the unknown accuracy of the open tree of life presents a risk of introducing error into phylogenetic models, as our ability to observe phylogenetic signal is partially dependent on the quality of a phylogeny. For studies aiming to use open tree of life topologies to account for correlative non-independence in the development of new finalised estimates of abundance change, this may be problematic and careful sensitivity analyses should be undertaken considering the open tree of life data as well as already established high quality phylogenies. However, our work is not focussed on deriving policy-shaping estimates of abundance change. Instead, we are solely focussed on showcasing the potential risk of ignoring correlative non-independence, in which case the greatest potential issue with using the lower quality open tree of life topologies is that our estimates of the phylogenetic signal, and in turn the variability around the average abundance trend, may be underestimated. Future work should explore the impact of phylogeny quality on inference in the abundance datasets, as a phylogeny is not available for most species, and so using topologies or taxonomies may be the only alternative.

We were only able to source a topology for 80% (N = 35,000) of species from the open tree of life; all other species were removed from the analysis. For studies with the overall aim of assessing biodiversity change, removing species could be problematic, as the overall trend would not be representative of all species. However, in our case, where the aim is to assess estimates gained from using different modelling approaches, trimming the data to species with an accompanying phylogeny has no impact on our conclusions. We then further trimmed the data to only include higher-quality time series, removing the following: time series that contained zeros (which we considered extreme cases of extinctions or recolonisations) and time series with missing abundance values for a given year throughout the sampling duration (i.e., we required consecutive abundance estimates.) We then further trimmed each dataset to only include time series which had greater than or equal to the median number of abundance observations i.e., including the longest 50% of time series in each dataset. These time series are characterised in Table S1. With our trimmed dataset, we derived a mean abundance in each year (in cases where there were more than one observation per year) for each time series, and then normalised each time series by dividing the maximum abundance in each time series by all other abundance values, thus constraining populations to a maximum of 1. After all of the above steps, we were left with 10 datasets, covering over 1,000 species (multiple time series per species) and 2,000 unique sites (multiple time series per unique coordinates), describing the trend in 26,000 populations (specific abundance time series) derived from more than 725,000 abundance estimates.

Modelling

Prior to designing our models, we first explored what models have been used in the literature to explore abundance change patterns. In this work, we focussed on studies trying to characterise the average change in abundances over time, rather than studies attempting to assess how many species are declining or increasing, as this avoids discretizing a numeric value i.e. by assessing the average change, we avoid having to define what change is necessary to be classified as a ‘decline’. Using a non-systematic and non-random search of the literature, we identified 18 studies/reports looking at abundance change, and we characterised each studies modelling approach (Table S2). Across the 18 approaches, there was no clear ‘standard’ approach utilised by all studies, and instead, there was a general pattern of each study adapting and utilising one of two broad methods:

1) Log-linear modelling - where the natural log of abundance is modelled against time, whilst controlling for some site/location level variance through either fixed or random effects. In this approach, an important decision appears to be whether the study allows abundance trends to vary across sites and species (e.g., random slopes) or holds them static so all species and sites have the same abundance trend but with differing mean abundances (e.g., random intercepts). This log-linear modelling approach, sometimes utilising poisson generalised linear models with count data, was present in 16 of the 18 studies.

2) Decomposition - where abundance time series were broken down into time series of rate changes (e.g., time series of lambdas), which are then aggregated at different spatial and taxonomic scales. This approach was less common, occurring in 2 of the 18 approaches.

**Table S2.** Description of different abundance change models used in the literature, and how this model matches to the models we test. Our descriptions purely represent an approximate summary of each model's core structure, and so some of our model descriptions may be incomplete. As such, we do draw any direct comparison between any model in this table and our newly developed correlated effect model.

| Approach | Model | Model match |
|---|---|---|
| 1 | For each species, a poisson generalised linear model is run where abundance is predicted by two categorical fixed effects of site and year. This assumes all sites have the same linear trend but varying intercept, and that abundance observations at neighbouring points in time are independent.<br><br>To produce national averages, the mean predicted abundance estimates (average across all sites) for each species are derived, and then abundance estimates across species are smoothed i.e., not acknowledging the variable trends in species, and simply following the general pattern in abundances<br><br>Reference: (*20*)<br><br>Notes: Used on the UK Breeding Bird Survey and Common Bird Survey | **Random intercept/ Random slope**: Model assumes all sites for any given species have the same trend (like model 1),), or that species have different trends (model 2). The trend-aggregation is unlike any of the models we use, as it ignores variable rates of change in species, and instead focuses on the average abundance across species in each year. |
| 2 | For each species, a poisson generalised additive model is run where abundance is predicted by a categorical fixed effects of site and a smoothing effect of year. This assumes all sites have the same smoothed (likely non-linear) trend.<br><br>To produce national averages, the mean predicted abundance estimates (average across all sites) for each species are derived, and then abundance estimates across species are smoothed i.e., not acknowledging the variable trends in species, and simply following the general pattern in abundances<br><br>Reference: (*21*)<br><br>Notes: Used on the UK Breeding Bird Survey and Common Bird Survey | **Random intercept/ Random slope**: As above |
| 3 | For each species, a poisson generalised additive model is run where abundance is predicted by a random slope of site interacting with a smoothing effect of year. This allows sites to have varying smoothed (likely non-linear) trends. | **Random slope but with temporal non-linearity of Correlated effect model**: Species and sites are allowed to have varying trends. Site |

| | | |
|---|---|---|
| | Focus is on the species level trends. Approach to aggregate trends is not described<br><br>Reference: (*22*)<br><br>Notes: Used on the North American Breeding Bird Survey | level trends are smooth, not linear. No trend aggregation described |
| 4 | For each species, a poisson generalised additive model is run where abundance is predicted by a random slope of site interacting with a smoothing effect of year. Unlike the above model, this also includes an additional random term to account for site-level random deviations from the smooth. This allows sites to have varying smoothed (likely non-linear) trends.<br><br>Focus is on the species level trends. Approach to aggregate trends is not described<br><br>Reference: (*22*)<br><br>Notes: Used on the North American Breeding Bird Survey | **Random slope but with temporal non-linearity of Correlated effect model**:<br>As above |
| 5 | For each species, a poisson generalised linear model is run where abundance is predicted by a random slope of site interacting with a linear effect of year. This model also includes an additional random term *c* to account for site-level random intercepts. This allows sites to have varying log-linear trends.<br><br>Focus is on the species level trends. Approach to aggregate trends is not described<br><br>Reference: (*8*)<br><br>Notes: Used on the North American Breeding Bird Survey | **Random slope**:<br>Species and sites are allowed to have varying trends. Site level trends are linear. No trend aggregation described. |
| 6 | Study calculates yearly lambdas (Nt/Nt+1) for each pair of abundance observations in every population time series, and then averages lambdas across all populations in each year to report the estimated mean lambda/year.<br><br>Study also uses a second method, calculating the proportion of populations increasing vs decreasing with kendall tau correlation between abundance and time<br><br>No explicit accounting for autocorrelation | **Random slope**: Only a tentative match - this approach completely ignores hierarchical structure in the data. |

| | | |
|---|---|---|
| | Reference: (*23*) | |
| 7 | Study derives mean rate of change (lambda) per population time series by regressing the natural logarithm of abundance against the continuous variable year.<br><br>Trends (rates of change) are then averaged to aggregate species/site level estimates<br><br>Reference: (*24*) | **Random slope**: Similar to the random slope approach, but derives population level slopes in a pre-modelling step, instead of within the model |
| 8 | The natural log of abundance (biomass) is regressed against year, with a random intercept term for each population. Model also captures temporal autocorrelation with an auto-regressive 1 process to indicate correlation between abundance neighbouring observations<br><br>References: (*25*) | **Random intercept but with the temporal term from the Correlated effect model**: Is similar in structure to the random intercept model, but without the taxonomic and site-level random effects. |
| 9 | For each species, a poisson generalised linear model is run, with abundance as the response, regressed against year (treated continuously) and site (as a factor). This assumes abundance-time trends are the same across all sites, but acknowledges that species will have varying trends.<br><br>There is no information on how trends are aggregated at the national/global-level<br><br>Reference: (*26*) | **Random intercept/Random slope**: The assumption of all a species sites having the same trend matches the random intercept model. But varying species trends matches the random slope |
| 10 | A poisson generalised linear model is run, with abundance as the response, regressed against year (treated continuously), with site and region as nested random intercepts. This assumes abundance-time trends are the same across all sites, but further, as all species are included in the one model, each site-level intercept represents multiple species. Species trends are ignored<br><br>Reference: (*9*)<br><br>Notes: Used on the German insects data set (*9*) | **Random intercept**:<br>All sites are assumed to have the same trend. Not an exact match to the random intercept model as species are not treated as a distinct unit (random intercept) |
| 11 | A poisson generalised linear model is run, with abundance as the response, regressed against year (treated continuously), with site and region as nested random intercepts. There is also an | **Random intercept**: As above |

| | | |
|---|---|---|
| | additional random intercept of year, so year as treated as both a fixed and random effect. This model assumes abundance-time trends are the same across all sites, but further, as all species are included in the one model, each site-level intercept represents multiple species. Species trends are ignored<br><br>Reference: (*27*)<br><br>Notes: Used on the German insects data set (*27*) | |
| 12 | A poisson generalised linear model is run, with abundance as the response, regressed against year (treated continuously). The model includes correlated random slopes of abundance varying by year differently in sites and regions. This model assumes abundance-time trends differ across sites. Further, as all species are included in the one model, each site-level intercept and slope represents multiple species. Species trends are ignored<br><br>Reference: (*27*)<br><br>Notes: Used on the German insects data set (*9*) | **Random slope**:<br>Sites are assumed to have varying trends. Not an exact match to the random slope model as species are not treated as a distinct units (random slopes) |
| 13 | Study derives mean rate of change (lambda) per population time series by regressing the natural logarithm of abundance against the continuous variable year.<br><br>Trends (rates of change) are then aggregatesaggregated to species/site level estimates in a further mixed model<br><br>Reference: (*28*)<br><br>Notes: Used on the BioTIME dataset (*10*) | **Random slope**: Most closely resembles the random slope model, but instead of calculating rates of change within the model (as in our random slope model), rates of change are estimated in a preliminary step. |
| 14 | Each abundance time series is smoothed with a generalised additive model, to produce predicted values of abundance. Yearly pairwise-lambdas are then derived from these predicted values of abundance, essentially decomposing abundance time series into rate change (lambda) time series.<br><br>Lambdas are averaged each year at the species-level. Species level lambdas are then averaged to produce a global level estimate.<br><br>Reference: (*29*) | **Decomposition model** |

| | | |
|---|---|---|
| | Notes: Used on a sample of the Living Planet Data (*5*) | |
| 15 | The natural log of abundance is regressed against an autoregressive-1 temporal term, grouped at the population level, which allows non-linear abundance change to occur over time. Random intercepts and slopes and used to handle the hierarchical structure in the data, and notably allows different sites and datasets to have different abundance trends. However, species level information is not used in the model, and so even at the finest resolution, each intercept/slope represents multiple species.<br><br>Reference: (*30*) | **Random slope with a temporal autocorrelation term**: Most closely resembles the random slope model, but handles and accounts for temporal autocorrelation between neighbouring abundance observations. |
| 16 | Study derives estimated rate of change per population time series from a mann-kendall correlation between abundance and time<br><br>Trends (rates of change) are then aggregates to species/site level estimates in a further meta regression<br><br>Reference: (*3*)<br><br>Notes: Used on the European insects dataset (*3*) | **Random slope**: Most closely resembles the random slope model, but instead of calculating rates of change within the model (as in our random slope model), rates of change are estimated in a preliminary step. |
| 17 | For each species, a poisson generalised linear model is run where abundance is predicted by two categorical fixed effects of site and year. This assumes all sites have the same linear trend but varying intercept, and that abundance observations at neighbouring points in time are independent.<br><br>Site-level estimates of abundance are combined to produce total counts for each species in each year. Approach for aggregating to national-level and global-level is unclear.<br><br>Reference: (*31*)<br><br>Notes: Used on the PECBMS dataset (*32*) | **Random intercept/ Random slope whilst handling temporal autocorrelation**: Model assumes all sites for any given species have the same trend (like model 1), but allows species to have different trends (model 2). The trend-aggregation is unlike any of the models we use, as it ignores variable rates of change in species, and instead focuses on the average abundance across species in each year. |
| 18 | Each abundance time series is smoothed with a generalised additive model, to produce predicted values of abundance. Yearly pairwise-lambdas are then derived from these predicted values of | **Decomposition model** |

| | |
|---|---|
| | abundance, essentially decomposing abundance time series into rate change (lambda) time series.<br><br>Lambdas are averaged each year at the species-level. Species level lambdas are then averaged to produce a global level estimate.<br><br>Reference: (*33*)<br><br>Notes: Used on a sample of the Living Planet Data (*5*) |

Given the decomposition modelling approach was comparably rare, we instead focus solely on the log-linear mixed modelling approaches, and develop the three models:

*Model 1. Random intercept*

In model 1, we fit a linear mixed effect model between the natural logarithm of normalised-abundance and year, with five random intercepts: population (the unique time series), site (unique locations), region (broader spatial category to nest sites; measured as the continent or ocean the site occurs in), species (unique species), and genus (broader taxonomic category to nest species; measured as the parent node to the species tip). Within the model, we do not specify any nesting between the site and species random intercepts as the hierarchical structure of the data is poorly defined e.g., whilst populations always occur within a species and site, some species are nested in sites, and some sites are nested in species, creating a crossed random effect design. Model 1 assumes all populations, sites, regions, species, and genera have the same trend in abundance.

$$u_{ijklm} = u_i^S + u_j^L + u_k^P + u_l^G + u_m^R$$

$$\boldsymbol{u^S} \sim N(0, \sigma^2{}_S I)$$

$$\boldsymbol{u^L} \sim N(0, \sigma^2{}_L I)$$

$$\boldsymbol{u^P} \sim N(0, \sigma^2{}_P I)$$

$$\boldsymbol{u^G} \sim N(0, \sigma^2{}_G I)$$

$$\boldsymbol{u^R} \sim N(0, \sigma^2{}_R I)$$

$$\bar{y}_{ijklmt} = a + [\beta]x_{ijklmt\ +} + u_{ijklm}$$

$$y_{ijklm} \quad N(\bar{y}_{ijklmt}, \sigma^2{}_E I)$$

Where $u$ represents the independent random intercept terms for species ($S$, index $i$), locations ($L$, index $j$), populations ($P$, index $k$), genera ($G$, index $l$) and regions ($R$, index $m$), all following a gaussian normal-distribution, with each varying according to their respective sigma hyperprior. These random intercepts vary around the overall model intercept ($a$), with a slope coefficient of β describing abundance change over years ($x$). This formula describes expected abundances ($\bar{y}$) for each intercept grouping, at time point $t$ (indexing of each abundance observation). $y$ represents a vector of normalised abundances for each population (index $k$), drawn from a gaussian normal-distribution with a mean $\bar{y}$ and a residual error of $\sigma^2_E$. $I$ describes the identity matrix of the error terms.

*Model 2. Random slope*

In model 2, we develop a linear mixed effect model, where we regress the natural logarithm of normalised-abundance against year, including population, site, region, species, and genus all as random slopes. This allows abundance-year slope coefficients to vary for each category in each random slope term (e.g., each species can have a different slope) - not simply differing intercepts as in model 1. To reduce model parameters, we centre the year and normalised abundances of each population time series at zero e.g. subtracting each year by the mean year in each population, and subtracting the log of each abundance value by the mean log abundance value in each population. This centering fixes the y and x intercepts at zero for each slope.

$$u_{ijklm} = u^S_i + u^L_j + u^P_k + u^G_l + u^R_m$$

$$\boldsymbol{u^S} \sim N(0,\sigma^2_S I)$$

$$\boldsymbol{u^L} \sim N(0,\sigma^2_L I)$$

$$\boldsymbol{u^P} \sim N(0,\sigma^2_P I)$$

$$\boldsymbol{u^G} \sim N(0,\sigma^2_G I)$$

$$\boldsymbol{u^R} \sim N(0,\sigma^2_R I)$$

$$\bar{y}_{ijklmt} = a + [\beta + u_{ijklm}]x_{ijklmt}$$

$$y_{ijklm} \quad N(\bar{y}_{ijklmt}, \sigma^2_E I)$$

Where $u$ represents the independent random slope terms for species ($S$, index $i$), locations ($L$, index $j$), populations ($P$, index $k$), genera ($G$, index $l$) and regions ($R$, index $m$), all following a gaussian normal-distribution, with each varying according to their respective sigma hyperprior. These independent random slopes vary around the overall slope coefficient of β describing abundance change over years ($x$) - meaning the abundance-time slope coefficient is allowed to vary in each species, location, population, genera, and region. $a$ describes the overall model intercept, which is included to support model convergence, but has a value of c.0 given the centering of the abundance and year values described above. This formula describes expected abundances ($\bar{y}$) for each slope grouping term, at time point $t$ (indexing of each abundance observation). $y$ represents a vector of normalised abundances for each population (index $k$), drawn from a gaussian normal-distribution with a mean $\bar{y}$ and a residual error of $\sigma^2{}_E$. $I$ describes the identity matrix of the error terms.

*Model 3. Correlated effect*

Model 3 is structurally similar to model 2, but accounts for correlative non-independence structures. For temporal non-independence, we model the population level time series with a discrete autoregressive-1 temporal process, which assumes neighbouring abundance observations within a time series will be more similar. To capture the spatial and phylogenetic correlative non-independence, we focus on non-independence across time series trends (instead of abundance observations), assuming trends in population abundances through time will be more similar in neighbouring sites and more closely related species. In model 1 and 2, we try to capture this non-independence with grouping categories (genus and region). However, in the correlated effect model, we more explicitly describe shared correlations between every species and site by specifying the covariance structure of our site and species random slopes. The site covariance matrix was derived by developing a matrix that describes the Euclidean distance between each site. We normalised this matrix between 0 and 1, with values close to 1 indicating neighbouring sites, whilst values approaching 0 indicate distant sites. We then converted this correlation matrix into a variance-covariance matrix. The species covariance matrix was derived by extracting the variance-covariance matrix directly from the species' phylogeny.

$$u_{ijklm} = u_i^S + u_j^L + u_k^P + u_l^G + u_m^R$$

$$\boldsymbol{u^S} \sim N(0,\sigma^2{}_S I)$$

$$\boldsymbol{u^L} \sim N(0,\sigma^2{}_L I)$$

$$\boldsymbol{u^P} \sim N(0,\sigma^2{}_P I)$$

$$\boldsymbol{u^G} \sim N(0,\sigma^2{}_G I)$$

$$\boldsymbol{u^R} \sim N(0,\sigma^2{}_R I)$$

$$v_{ij} = v^{S}_{i} + v^{L}_{j}$$

$$\boldsymbol{v^{S}} \sim N(0,\sigma^{2}_{S}\Omega)$$

$$\boldsymbol{v^{L}} \sim N(0,\sigma^{2}_{L}\Delta)$$

$$\bar{y}_{ijkt} = a + [\beta + u_{ijklm} + v_{ij}]x_{ijkt}$$

$$y_{ijk} \quad N(\bar{y}_{ijkt}, \sigma^{2}_{E}I + \sigma^{2}_{A}\Theta)$$

Where $u$ represents the independent random slope terms for species ($S$, index $i$), locations ($L$, index $j$), populations ($P$, index $k$), genera ($G$, index $l$) and regions ($R$, index $m$), whilst $v$ represents the correlated random slope terms for species ($S$, index $i$) and locations ($L$, index $j$). All random slopes, independent and correlated, follow a gaussian normal-distribution, with each varying according to their respective sigma hyperprior. However, the independent and correlated slopes differ, as $u$ varies according to the identity matrix $I$, whilst $v$ varies according to variance-covariances $\Omega$ and $\Delta$ which specify that covariance is present in neighbouring sites and more closely related species. These independent and correlated random slopes vary around the overall slope coefficient of β describing abundance change over years ($x$) - meaning the abundance-time slope coefficient is allowed to vary in each species, location, and population. $a$ describes the overall model intercept. This formula describes expected abundances ($\bar{y}$) for each slope grouping term, at time point $t$ (indexing of each abundance observation). $y$ represents a vector of normalised abundances for each population (index $k$), drawn from a gaussian normal-distribution with a mean of $\bar{y}$. However, unlike model 2, the error term of this distribution has two components, the residual error of $\sigma^2_E$ as in model 2, and a new error term $\sigma^2_A\Theta$ which captures temporal non-independence by parameterising the correlation between neighbouring abundance values ($\Theta$, often called *rho*) and the left-over error from this process ($\sigma^2_A$).

Priors

Across all three models, we set vague normal priors on the fixed effects ($\beta_0$ - $\beta_1$), centered at zero, with a high standard deviation of 100. All random intercepts and slopes were assigned a normal prior, centered at zero, with an improper uniform hyperprior determining the standard deviation of normal priors. We used a uniform hyperprior following established recommendations (*34*). All models were run in INLA (*35*)

Outputs

We compare our three models across 3 key outputs:

*Measuring non-independence*

In our *Correlated effect* model we measure the presence of total non-independence as the proportion of variance captured by the combination of independent and correlative terms (i.e. random effects) for each component (e.g., temporal components), divided by the sum of the variance for all terms. Next, we assess if correlative terms are the larger contributor to this total non-independence, by dividing the proportion of variance captured by the correlated slopes, by the combination of the variance captured by the correlated and independent slopes (*h*). This was done separately for the spatial and phylogenetic terms. As the spatial and phylogenetic components each contains three terms (an independence species/location slope, an independent genera/region slope, and a correlated species slope), a h-value of 0.33 would indicate that the correlative slope captures an equal proportion of variance compared to the two independent slopes. A *h* greater than 0.33 indicates that correlative slopes account for more variation than independent random slopes. We measure temporal non-independence as the degree of correlation between neighbouring abundances (*rho).*

*Differing inference between the models*

Using the mean and 50% credible intervals of the global trend (overall abundance-time coefficients), we display abundance projections for each model in each dataset. These projections are based on an arbitrary baseline abundance of 100, set at the first year of available data in each dataset, and this abundance would change according to the overall coefficients and credible intervals. For instance, with a 1% annual rate of change, an abundance in year zero of 100, would become 101 in year 1, and 164 in year 50. The purpose of these projections is to showcase varying abundance trajectories under different model complexities.

Next, we note the number of the datasets where inference reverses (e.g., the global trend reverses direction from positive to negative, or remains consistent), and where uncertainty increases (the variance around the global trend is greater or smaller), comparing the random intercept and random slope models to the correlated effect model. To support these comparisons, we also report the absolute difference between the global trend of the correlated effect model, relative to the random intercept and random slope models. We modelled these differences in a linear model, with absolute difference as the response on the natural log scale, and model comparison as a factor (correlated effect versus random intercept, and correlated effect versus random slope). Further, we modelled the fold change in the standard deviation of the correlated effect model, relative to the other models, with fold change as the response on the natural log scale, and model comparison as a factor - see Figure S2 for marginal effects.

*Predictive performance*

We assess the predictive performance of the different models by determining their ability to predict final observations in time series', and their ability to predict population trends of a given species in a given location, both using the BioTIME dataset (*10*) as a case study. To test the predictive accuracy for the final observation in the time series, we removed the final observation half of the time-series (N = 3520) and predicted the missing values using each of the three models. We then compared the predicted value with the observed abundance value, recording the absolute error (the absolute difference between the predicted and observed value), and relationship ($\beta$) between the true (response) and predicted values from a linear model. To test the accuracy of the population prediction, we removed 10% (N = 704) of the populations from the dataset, and used the random slope and correlated effect model to estimate these missing population's trend coefficients, relative to population trend coefficients from the correlated effect model with no missing data (observed values). We measured performance using the same approaches as above, recording the absolute error and relationship ($\beta$) between the predicted and observed values. In the random slope model, the population trend coefficients were derived by adding the species, location, genus and region coefficients together, meaning missing population values can still be informed by other hierarchical information. For the correlated effect model, the species coefficient is informed by the phylogenetic variance-covariance matrix, as well as all hierarchical information in the random slope model.

*Phylogenetic and spatial distribution of abundance change*

To plot abundance change across a phylogeny, we extracted and plotted species rate of change coefficients from the correlated effect model and plotted these on the phylogeny associated with the BioTIME dataset. These species rate of change coefficients solely describe the genetic species terms (correlated slopes for species from the variance-covariance matrix) and not the phenotypic terms (independent slopes as in the random slope model). We solely show the genetic term as there was a high phylogenetic signal in BioTIME (h = 0.88).

To plot abundance change over space, we expanded the BioTIME spatial Euclidean distance matrix (describing distances between each time series) by supplementing it with a regularly gridded extent covering North America. This new grid had a latitudinal range of 15 to 70 and 1 degree spacing (e.g. 15, 16, etc.), and longitudinal range of -140 to -50 with 1 degree spacing. This new matrix allows us to estimate expected covariance (similarity) in abundance trends for any pair of 1 degree cells across North America. We then re-ran the BioTIME model with the new distance matrix and extracted the annual rate of change (%) in abundance from every cell across the extent. We only showcase the abundance trends across North America where abundance patterns were particularly stark. WithinWithin this North America region, we down-scaled the resolution from 1 degree to 0.1 degree using exponential interpolation.

### Supplementary results

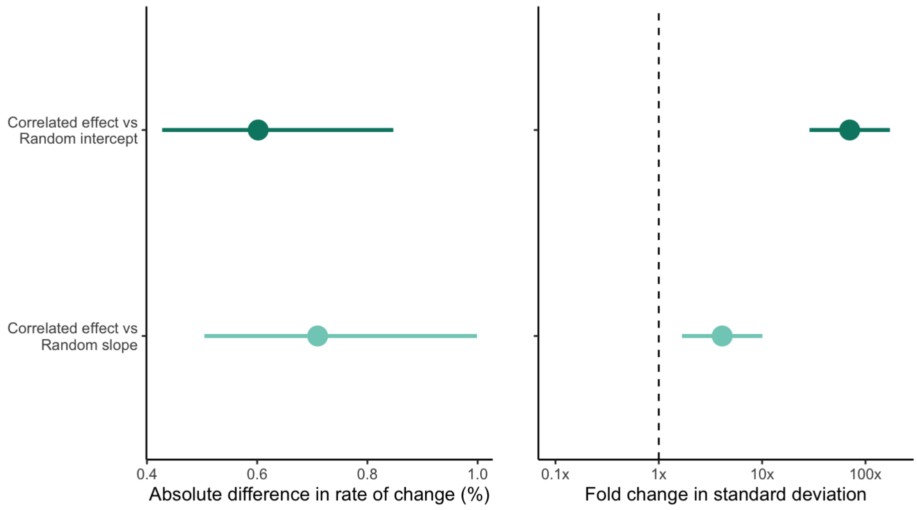

**Figure S2.** Absolute difference in the mean abundance-time coefficient (left) and fold-change in the standard deviation around said coefficient (right), comparing the correlated effect model to the random intercepts (dark green) and random slopes (light green) model. The x-axis on the fold-change figure is log-10 scaled, where a value of 1 would indicate the compared models have similar standard deviations. Values to the right of the line would indicate the standard deviation is X times greater in the correlated effect model.

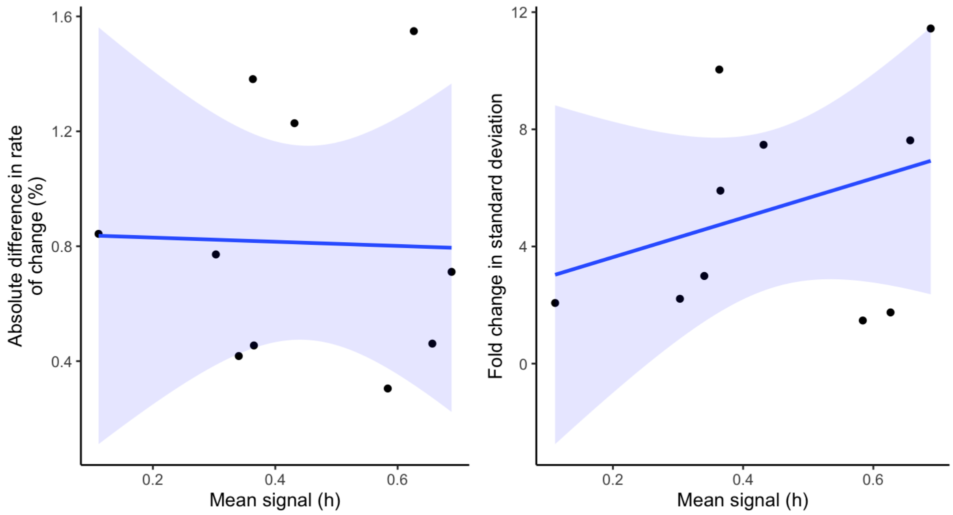

**Figure S3.** Impact of phylogenetic and spatial signal on inference. Left) Absolute difference in the abundance-time coefficient between the correlated effect and random slopes model, plotted against the mean signal (the mean of the phylogenetic and spatial signal). Right) Fold change in the standard deviation of the abundance-time coefficient between the correlated effect and random slopes model, plotted against the mean signal. Mean signal is calculated by finding the mean of the phylogenetic signal (variance captured by the phylogeny divided by the sum of the phylogenetic and non-phylogenetic variance) and spatial signal (as in the phylogeny). Each data point represents a dataset. Shading represents 95% confidence intervals from a linear model.

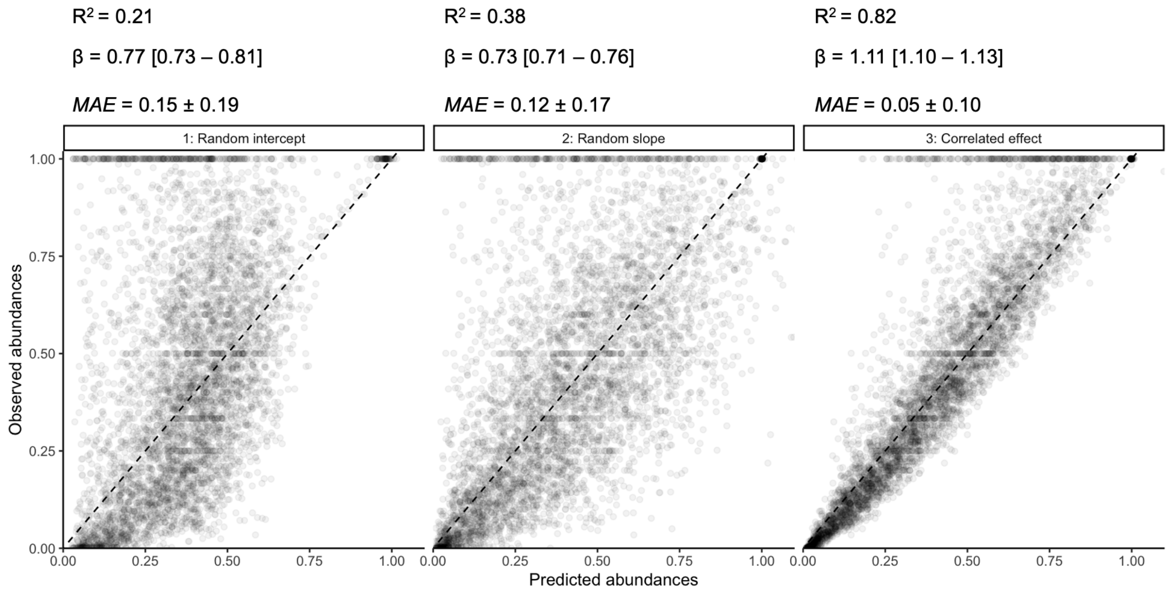

**Figure S4.** Ability of the three models to predict the missing-final abundance observation from half of the BioTIME time series, with the observed (the value we have removed) and predicted abundance value on the y and x axes, respectively. Points more closely following the diagonal dashed line exhibit a better fit. For each model we describe the R2 of the observed regressed against the predicted values, the beta slope coefficient from this model, and the median absolute error between the observed and predicted values (MAE).

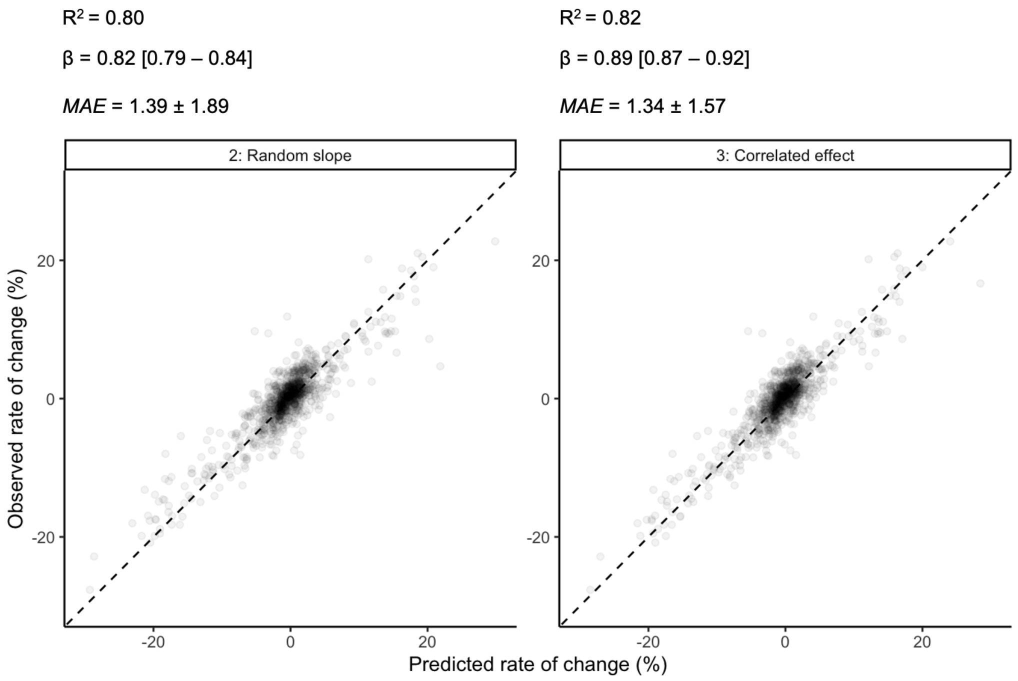

**Figure S5.** Ability of the random slope and correlated effect models to predict the 10% subset of population trends we removed from the BioTIME dataset, with the observed (the value we have removed) and predicted trend value on the y and x axes, respectively. Points more closely following the diagonal dashed line exhibit a better fit. For each model we describe the R2 of the observed regressed against the predicted values, the beta slope coefficient from this model, and the median absolute error between the observed and predicted values (MAE). We do not assess the ability of the random intercept model which assumes all populations have the same trend.

671–674 (2019).
10. M. Dornelas, L. H. Antão, F. Moyes, A. E. Bates, A. E. Magurran, D. Adam, A. A. Akhmetzhanova, W. Appeltans, J. M. Arcos, H. Arnold, N. Ayyappan, G. Badihi, A. H. Baird, M. Barbosa, T. E. Barreto, C. Bässler, A. Bellgrove, J. Belmaker, L. Benedetti-Cecchi, B. J. Bett, A. D. Bjorkman, M. Błażewicz, S. A. Blowes, C. P. Bloch, T. C. Bonebrake, S. Boyd, M. Bradford, A. J. Brooks, J. H. Brown, H. Bruelheide, P. Budy, F. Carvalho, E. Castañeda-Moya, C. A. Chen, J. F. Chamblee, T. J. Chase, L. Siegwart Collier, S. K. Collinge, R. Condit, E. J. Cooper, J. H. C. Cornelissen, U. Cotano, S. Kyle Crow, G. Damasceno, C. H. Davies, R. A. Davis, F. P. Day, S. Degraer, T. S. Doherty, T. E. Dunn, G. Durigan, J. E. Duffy, D. Edelist, G. J. Edgar, R. Elahi, S. C. Elmendorf, A. Enemar, S. K. M. Ernest, R. Escribano, M. Estiarte, B. S. Evans, T. Y. Fan, F. Turini Farah, L. Loureiro Fernandes, F. Z. Farneda, A. Fidelis, R. Fitt, A. M. Fosaa, G. A. Daher Correa Franco, G. E. Frank, W. R. Fraser, H. García, R. Cazzolla Gatti, O. Givan, E. Gorgone-Barbosa, W. A. Gould, C. Gries, G. D. Grossman, J. R. Gutierréz, S. Hale, M. E. Harmon, J. Harte, G. Haskins, D. L. Henshaw, L. Hermanutz, P. Hidalgo, P. Higuchi, A. Hoey, G. Van Hoey, A. Hofgaard, K. Holeck, R. D. Hollister, R. Holmes, M. Hoogenboom, C. hao Hsieh, S. P. Hubbell, F. Huettmann, C. L. Huffard, A. H. Hurlbert, N. Macedo Ivanauskas, D. Janík, U. Jandt, A. Jażdżewska, T. Johannessen, J. Johnstone, J. Jones, F. A. M. Jones, J. Kang, T. Kartawijaya, E. C. Keeley, D. A. Kelt, R. Kinnear, K. Klanderud, H. Knutsen, C. C. Koenig, A. R. Kortz, K. Král, L. A. Kuhnz, C. Y. Kuo, D. J. Kushner, C. Laguionie-Marchais, L. T. Lancaster, C. Min Lee, J. S. Lefcheck, E. Lévesque, D. Lightfoot, F. Lloret, J. D. Lloyd, A. López-Baucells, M. Louzao, J. S. Madin, B. Magnússon, S. Malamud, I. Matthews, K. P. McFarland, B. McGill, D. McKnight, W. O. McLarney, J. Meador, P. L. Meserve, D. J. Metcalfe, C. F. J. Meyer, A. Michelsen, N. Milchakova, T. Moens, E. Moland, J. Moore, C. Mathias Moreira, J. Müller, G. Murphy, I. H. Myers-Smith, R. W. Myster, A. Naumov, F. Neat, J. A. Nelson, M. Paul Nelson, S. F. Newton, N. Norden, J. C. Oliver, E. M. Olsen, V. G. Onipchenko, K. Pabis, R. J. Pabst, A. Paquette, S. Pardede, D. M. Paterson, R. Pélissier, J. Peñuelas, A. Pérez-Matus, O. Pizarro, F. Pomati, E. Post, H. H. T. Prins, J. C. Priscu, P. Provoost, K. L. Prudic, E. Pulliainen, B. R. Ramesh, O. Mendivil Ramos, A. Rassweiler, J. E. Rebelo, D. C. Reed, P. B. Reich, S. M. Remillard, A. J. Richardson, J. P. Richardson, I. van Rijn, R. Rocha, V. H. Rivera-Monroy, C. Rixen, K. P. Robinson, R. Ribeiro Rodrigues, D. de Cerqueira Rossa-Feres, L. Rudstam, H. Ruhl, C. S. Ruz, E. M. Sampaio, N. Rybicki, A. Rypel, S. Sal, B. Salgado, F. A. M. Santos, A. P. Savassi-Coutinho, S. Scanga, J. Schmidt, R. Schooley, F. Setiawan, K. T. Shao, G. R. Shaver, S. Sherman, T. W. Sherry, J. Siciński, C. Sievers, A. C. da Silva, F. Rodrigues da Silva, F. L. Silveira, J. Slingsby, T. Smart, S. J. Snell, N. A. Soudzilovskaia, G. B. G. Souza, F. Maluf Souza, V. Castro Souza, C. D. Stallings, R. Stanforth, E. H. Stanley, J. Mauro Sterza, M. Stevens, R. Stuart-Smith, Y. Rondon Suarez, S. Supp, J. Yoshio Tamashiro, S. Tarigan, G. P. Thiede, S. Thorn, A. Tolvanen, M. Teresa Zugliani Toniato, Ø. Totland, R. R. Twilley, G. Vaitkus, N. Valdivia, M. I. Vallejo, T. J. Valone, C. Van Colen, J. Vanaverbeke, F. Venturoli, H. M. Verheye, M. Vianna, R. P. Vieira, T. Vrška, C. Quang Vu, L. Van Vu, R. B. Waide, C. Waldock, D. Watts, S. Webb, T. Wesołowski, E. P. White, C. E. Widdicombe, D. Wilgers, R. Williams, S. B. Williams, M. Williamson, M. R. Willig, T. J. Willis, S. Wipf, K. D. Woods, E. J. Woehler, K. Zawada, M. L. Zettler, BioTIME: A database of biodiversity time series for the Anthropocene. *Glob. Ecol. Biogeogr.* **27**, 760–786 (2018).
11. S. P. R. Greenstreet, M. Moriarty, Manual for Version 3 of the Groundfish Survey Monitoring and Assessment Data Product. *Scott. Mar. Freshw. Sci.* **8** (2017).
12. T. F. Johnson, P. Cruz, N. J. B. Isaac, A. Paviolo, M. González-Suárez, CaPTrends: A database of large carnivoran population trends from around the world. *Glob. Ecol. Biogeogr.* **n/a** (2022), doi:10.1111/geb.13587.
13. F. Michonneau, J. W. Brown, D. J. Winter, rotl: an R package to interact with the Open Tree of Life data. *Methods Ecol. Evol.* **7**, 1476–1481 (2016).
14. OpenTree, Open Tree of Life Synthetic Tree version 13.4, (available at

https://doi.org/10.5281/zenodo.3937741).
15. R. M. Fisher, L. M. Henry, C. K. Cornwallis, E. T. Kiers, S. A. West, The evolution of host-symbiont dependence. *Nat. Commun.* **8**, 15973 (2017).
16. R. A. Morais, D. R. Bellwood, Global drivers of reef fish growth. *Fish Fish.* **19**, 874–889 (2018).
17. E. Hamann, C. Blevins, S. J. Franks, M. I. Jameel, J. T. Anderson, Climate change alters plant–herbivore interactions. *New Phytol.* **229**, 1894–1910 (2021).
18. A. Grafen, W. D. Hamilton, The phylogenetic regression. *Philos. Trans. R. Soc. Lond. B Biol. Sci.* **326**, 119–157 (1989).
19. E. Paradis, K. Schliep, ape 5.0: an environment for modern phylogenetics and evolutionary analyses in R. *Bioinformatics*. **35**, 526–528 (2019).
20. BTO, Breeding Bird Survey (2020), (available at https://www.bto.org/our-science/publications/birdtrends/2020/methods/breeding-bird-survey).
21. R. M. Fewster, S. T. Buckland, G. M. Siriwardena, S. R. Baillie, J. D. Wilson, Analysis of Population Trends for Farmland Birds Using Generalized Additive Models. *Ecology*. **81**, 1970–1984 (2000).
22. A. C. Smith, B. P. M. Edwards, North American Breeding Bird Survey status and trend estimates to inform a wide range of conservation needs, using a flexible Bayesian hierarchical generalized additive model. *Ornithol. Appl.* **123**, duaa065 (2021).
23. J. E. Houlahan, C. S. Findlay, B. R. Schmidt, A. H. Meyer, S. L. Kuzmin, Quantitative evidence for global amphibian population declines. *Nature*. **404**, 752–755 (2000).
24. C. S. Robbins, J. R. Sauer, R. S. Greenberg, S. Droege, Population declines in North American birds that migrate to the neotropics. *Proc. Natl. Acad. Sci.* **86**, 7658–7662 (1989).
25. J. A. Hutchings, C. Minto, D. Ricard, J. K. Baum, O. P. Jensen, Trends in the abundance of marine fishes. *Can. J. Fish. Aquat. Sci.* **67**, 1205–1210 (2010).
26. N. Poulet, L. Beaulaton, S. Dembski, Time trends in fish populations in metropolitan France: insights from national monitoring data. *J. Fish Biol.* **79**, 1436–1452 (2011).
27. G. N. Daskalova, A. B. Phillimore, I. H. Myers-Smith, Accounting for year effects and sampling error in temporal analyses of invertebrate population and biodiversity change: a comment on Seibold et al. 2019. *Insect Conserv. Divers.* **14**, 149–154 (2021).
28. M. Dornelas, N. J. Gotelli, H. Shimadzu, F. Moyes, A. E. Magurran, B. J. McGill, A balance of winners and losers in the Anthropocene. *Ecol. Lett.* (2019), doi:10.1111/ele.13242.
29. F. He, C. Zarfl, V. Bremerich, J. N. W. David, Z. Hogan, G. Kalinkat, K. Tockner, S. C. Jähnig, The global decline of freshwater megafauna. *Glob. Change Biol.* **25**, 3883–3892 (2019).
30. R. van Klink, D. E. Bowler, K. B. Gongalsky, A. B. Swengel, A. Gentile, J. M. Chase, Meta-analysis reveals declines in terrestrial but increases in freshwater insect abundances. *Science* (2020), doi:10.1126/science.aax9931.
31. PECBMS, Production of national indices and trends (2022), (available at https://pecbms.info/methods/pecbms-methods/1-national-species-indices-and-trends/1-2-production-of-national-indices-and-trends/).
32. PECBMS, Pan-European Common Bird Monitoring Schem (2021), (available at https://pecbms.info/).
33. L. McRae, S. Deinet, R. Freeman, The Diversity-Weighted Living Planet Index: Controlling for Taxonomic Bias in a Global Biodiversity Indicator. *PLOS ONE*. **12**, e0169156 (2017).
34. A. Gelman, Prior Distribution for Variance Parameters in Hierarchical Models. *Bayesian Anal.* (2006).
35. H. Rue, S. Martino, N. Chopin, Approximate Bayesian inference for latent Gaussian models by using integrated nested Laplace approximations. *J. R. Stat. Soc. Ser. B Stat. Methodol.* **71**, 319–392 (2009).
